# Automated auditory brainstem response peak estimation using a convolutional neural net

**DOI:** 10.64898/2026.06.30.735643

**Authors:** Jax P. Marrone, Meredith C. Ziliak, Edward L. Bartlett

## Abstract

Auditory brainstem responses (ABRs) are a core part of objective functional evaluations of hearing sensitivity and subcortical auditory transmission. Manual assessments of ABR waveforms are still a primary means by which thresholds and peak amplitudes and latencies are measured, which is time-consuming and prone to user variability. Automated methods have offered promising alternatives for ABR classification, but they have sometimes been limited in accuracy or robustness. Here, we developed and tested a supervised convolutional neural network (CNN) based ABR peak classifier that works across sound levels and sound frequencies that can be run quickly on a personal computer using single or dual-channel ABR inputs. For ABR peaks I, III, IV, and V, the classifier achieved over 95% accuracy. High accuracy was maintained even after noise-exposure causing temporary or permanent threshold shifts, and over 90% of peaks were within 0.041 ms (1 sample) of the manually identified peak. Only a few hundred samples were needed to train the network, making it widely amenable to smaller data studies or where the number of subjects or sessions may be low.

**Highlights:**

- Supervised CNN automatic ABR peak classifier built for 1-2 channel inputs
- Accuracy for peaks was at least 95% for all peaks
- Accuracy was robust across sound levels and noise-exposure
- A relatively small number of labeled samples were sufficient for training

**CRediT author contributions:** JPM: Conceptualization, Methodology, Investigation, Formal analysis, Writing - original draft, Writing - review & editing. MCZ: Methodology, Investigation. ELB: Conceptualization, Methodology, Resources, Supervision, Writing – original draft, review & editing, Funding acquisition.

## Introduction

Auditory brainstem responses (ABRs) are a fundamental electrophysiological measure of auditory pathway function (Hecox and Galambos 1974, Starr and Hamilton 1976). In addition to their use in clinical diagnostics, ABRs are widely applied to investigate auditory processing changes across damage paradigms, such as aging or noise exposure, in humans and preclinical animal models (Sergeyenko et al. 2013, Parthasarathy and Kujawa 2018, Lai et al. 2017, Race et al. 2017). ABR thresholds are considered the objective audiologic standard for hearing function. However, the amplitudes and latencies of ABR waveforms are rich sources of additional information that track propagation through regions of the ascending auditory pathway (Moller et al. 1983, Ponton et al. 1996, Bidelman 2015).

Obtaining these waveform measurements often relies on arduous, manual peak-picking of wave-specific peaks and troughs. Standard audiogram acquisition requires presenting stimuli across multiple sound levels for each test frequency, with each level-frequency combination needing separate labeling. In multi-subject longitudinal studies, these demands make manual peak picking prohibitively time-consuming. Labeling is further complicated by ambiguous morphologies, often induced by acoustic trauma or mutations (Ingham et al. 2011), including split peaks and poorly demarcated troughs, which diminish both inter- and intra-reviewer consistency (Vidler and Parkert 2004). Automated peak identification is therefore critical to improve the efficiency and reproducibility of ABR analysis.

Several approaches have attempted to address this problem. Derivative-based detectors marked peaks at the first-derivative zero crossings of the ABR signal (Fridman et al. 1982, Bradley and Wilson 2005). Although accurate for high-quality recordings, this procedure hinged on rigid, wave-specific latency windows that fail when response timing is altered by auditory damage. Template matching methods aligned full ABR traces to a labeled reference via cross-correlation and transferred peak locations from reference to observation (Elberling 1979). Alignment was anchored at wave V, implicitly fixing inter-peak intervals and minimizing both normative variability and the larger deviations introduced by auditory injury. Employing wave-specific, individually matched templates partially alleviated this issue, although accuracy remained contingent on fidelity to an idealized morphology (Vannier et al. 2002, Valderrama et al. 2014). Dynamic time warping (DTW) offered a less stringent alignment procedure by applying non-linear temporal transformations to fit local regions of the ABR to a reference trace. Early adoptions discretely warped each candidate waveform to a predefined target signal, risking bias when the template was unrepresentative (Picton et al. 1988). A subsequent variant matched ABRs to a data-derived structural average using smooth, monotonic warps, improving generalizability. The smoothness assumption, however, was susceptible to failure on noisy data, obscuring irregular waveform distortions (Krumbholz et al. 2020). Gaussian mixture model feature extraction techniques (GMM-FET) replaced temporal warping with parametric wave decomposition, modeling the ABR as a superposition of overlapping Gaussian components (Kamerer 2024). This method explicitly accommodated inter-wave interactions without dependence on a fixed template. However, it assumed approximately Gaussian wave shapes and has been validated so far only at high stimulus intensities (100-110 dB SPL); performance at lower levels remains untested.

These approaches share two central limitations: reliance on explicit morphological priors and limited robustness for noisy, near-threshold, or pathologically altered responses. Neural networks offer a more flexible alternative, but published implementations remain constrained. Early multilayer perceptrons demonstrated feasibility but were limited to click-evoked ABRs and did not report exact-match accuracy (Tian et al. 1997); bidirectional long short-term memory (BiLSTM) networks achieved ∼92% accuracy, but only under a liberal temporal tolerance (±0.2 ms) (Chen et al. 2020); and convolutional recurrent neural networks (CRNNs) reported ∼96% accuracy within a ±0.1 ms tolerance, but were evaluated only in normal-hearing cohorts (McKearney et al. 2025). Collectively, these studies were evaluated under relatively restricted conditions, with limited assessment across diverse ABR stimuli and types and severities of auditory damage.

Convolutional neural networks (CNNs) emerge as a natural continuation for automated peak picking. CNNs are well suited to biomedical signal processing, as their convolutional filters capture both local temporal and spatial features, encoding important electrophysiological properties (Rodriguez et al. 2023). Compared with fully connected or recurrent architectures, they balance accuracy with computational efficiency while scaling well to large, heterogeneous datasets (Alzubaidi et al. 2021, Lee et al. 2024). Their ability to learn directly from diverse input data allows CNNs to accommodate variable ABR morphologies without enforcing fixed templates or latency windows, enabling robust analysis (Lee et al. 2024).

This study introduces a CNN classifier for two-channel ABRs acquired using click and tone-burst stimuli (4, 8, 10, and 16 kHz). The model accurately identified waves I and III from a midline electrode (channel 1) and waves I, IV, and V from an interaural electrode (channel 2) (Parthasarathy and Bartlett 2012). Coupling the initial predictions with a local extrema adjustment refined the estimated time and amplitude of each target, ensuring exact coordinate matching rather than tolerance-based scoring. To capture a broad range of ABR morphologies, the model was evaluated across both normal and impaired, noise-exposed responses.

Two-channel ABR recordings captured both threshold elevations and waveform alterations caused by auditory damage (Parthasarathy and Bartlett 2012). Prior work has documented persistent threshold increases following mild blast exposure, natural aging, and small arms fire (SAF) like noise exposure (Parthasarathy et al. 2014, Altschuler et al. 2019, Han et al. 2021). Across these paradigms, varying changes in ABR wave amplitudes were observed, suggesting that while threshold shifts are a common outcome, the underlying morphological and recovery patterns may differ. The CNN approach discussed enables rapid, accurate waveform analysis, offering the potential to define these distinctions and thereby advance understanding of auditory injury mechanisms.

## Methods

### Subjects

Eighty-one Fischer 344 rats (3-7 months old) were used in this study. Of these, 73 animals (36 males, 37 females) were exposed to acute high-intensity noise simulating small arms fire (SAF), and 8 animals (4 males, 4 females) were exposed to continuous moderate noise. Detailed group assignments are provided in **Error! Reference source not found.**. All animals were kept and raised in standard laboratory animal housing conditions. All procedures were approved by the Purdue Animal Care and Use Committee (PACUC Protocol Nos. 2105002151 and 1204000631) and conducted in accordance with institutional and federal guidelines for animal research.

### SAF Exposure

The SAF exposure stimulus comprised a sequence of 12 biphasic pulses (150 μs/phase) repeated 50 times over 2.5 minutes. Peak sound pressure levels (SPLs) varied by group and were 80 (sham), 131, 134, 137.0, and 140 dB SPL. dB are listed as dB pSPL.

Anesthesia was administered intramuscularly using a ketamine/dexmedetomidine (Dexdomitor) cocktail. Sixty-nine animals received ketamine 60 mg/kg and dexmedetomidine 0.1 mg/kg; four received ketamine 40 mg/kg with the same dexmedetomidine dose. Within the 60 mg/kg group, 19 rats were exposed to 80 dB SPL (sham group), 2 to 131 dB, 27 to 134 dB, 11 to 137 dB, and 10 to 140 dB. All rats in the 40 mg/kg group were exposed to 134 dB SPL. Subjects were classified as functional, meaning a functional threshold shift, if they exhibited a ≥10 dB threshold shift 7 days post-insult.

Exposures were delivered from a JBL Selenium D3500Ti-Nd speaker positioned 5 in above the head, with animals centered beneath the acoustic axis. Rats were placed on a platform inside a 34 x 19.5 x 27.5 in (L x W x H) chamber constructed from 0.5-in compressed wood, topped with a matching lid containing a centered, 3-in-diameter cutout for the speaker. The assembly was housed in a 9 x 9 ft double-walled acoustic chamber (Industrial Acoustics Corporation).

### Continuous Exposure

Subjects in the continuous exposure cohort were exposed to constant pink noise at 87.5 dB SPL (A-weighted) for 40 hours per week over a 4-week period (4 days/week, 10 hours/day). Animals remained in their home cages with ad libitum access to food and water. Two cages were placed at a time inside the same exposure chamber used for the SAF condition.

### ABR Recordings

Two-channel auditory brainstem response (ABR) recordings were conducted at baseline (pre-exposure) and at 7, 14, 28, and 56 days post-exposure for SAF subjects, and at baseline and weekly intervals (1-4 weeks) for continuous noise subjects.

Animals were anesthetized briefly with 1.5-3% isoflurane. Subdermal needle electrodes (Ambu) were inserted in the following configuration: the channel 1 positive electrode along the midline of the head (midsagittal, oriented Fz-Cz), the channel 2 positive electrode along the interaural line (C3-C4), the negative electrode beneath the mastoid of the right ear ipsilateral to the speaker, and the ground electrode in the lower back (Fig. ***1***A). This placement followed configurations established in previous studies (Parthasarathy and Bartlett 2012, Lai et al. 2017). Electrode impedance was verified to be <1 kΩ using a low-impedance amplifier (RA4LI, Tucker-Davis Technologies).

**Fig. 1:**
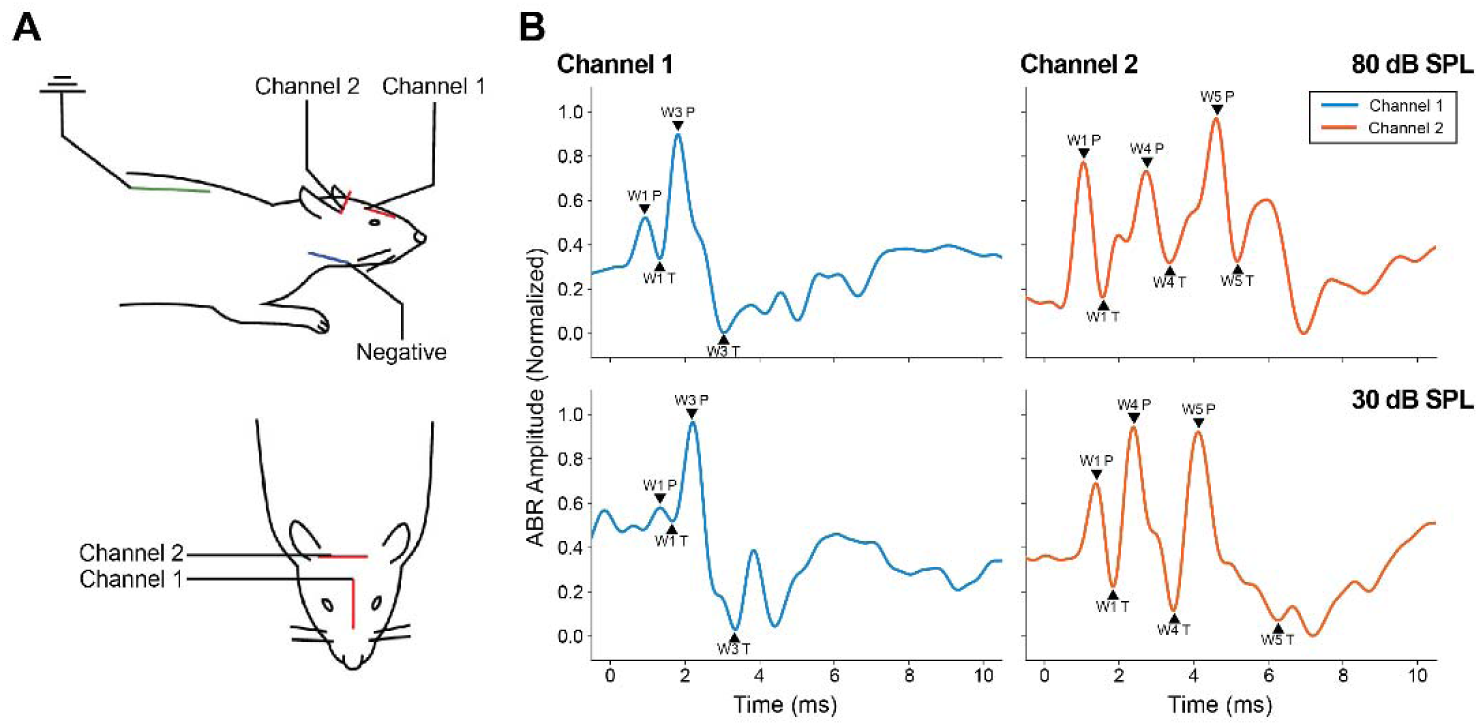
Two-channel electrode placement for auditory brainstem response (ABR) recordings (A) and representative ABR waveforms at high (80 dB) and low (30 dB) stimulus levels (B). ABRs were treated as discrete signals, with the index positions of each peak and trough (black triangles) manually labeled as ground truth targets for CNN training. Signals were filtered to 80-1500 Hz, amplitude-normalized to 0-1, and truncated to frequency-specific windows of equal duration (10.240 ms), each including a 3.2 ms free field delay. Ch1 shows distinct waves I and III, and Ch2 shows distinct waves I, IV, and V.

After electrode placement, animals were sedated with an intramuscular injection of dexmedetomidine (0.15-0.25 mg/kg). Recordings were initiated at least 11 minutes after isoflurane removal to minimize residual anesthetic effects. The animals could respond to pain and acoustic stimuli, but sat calmly under sedation, allowing 4 hours of recording time.

Acoustic stimuli were delivered free field to the right ear (90° azimuth, 0° azimuth reference directly in front of the animal’s face) using a calibrated Bowers & Wilkins speaker positioned 115 cm from the animal. ABRs were evoked using alternating polarity rectangular clicks (0.033 ms duration) and tone bursts (2 ms duration, 0.5 ms cos^2^ rise/fall time; frequencies 4, 8, 10, and 16 kHz), each with a 10 ms acquisition window, averaged over 1,500 repetitions. Stimuli were presented in 5 dB increments from 5 to 80 dB SPL (8, 10, and 16 kHz) and from 10 to 80 dB SPL (click and 4 kHz).

### Data Preprocessing

ABR signals were bandpass filtered using an 80 Hz high-pass and a 1,500 Hz low-pass filter. Only above-threshold ABR responses were included for CNN training. Thresholds were visually defined as the minimum stimulus level that produced a distinct ABR waveform. Signals were amplitude-normalized to 0-1 and truncated to equal-duration frequency-specific windows (10.240 ms), with start times defined relative to trigger onset (click, 3.686 ms; 4 kHz, 4.506 ms; 8 kHz, 4.178 ms; 10 kHz, 4.137 ms; 16 kHz, 4.014 ms).

A total of 2,898 ABRs were manually labeled for Ch1 and 2,788 for Ch2 (Table ***2***). For pure-tone ABRs, a random subset of recordings per frequency was selected for annotation. Recordings identified as below threshold were excluded, resulting in variation in sample size across stimuli and differences in the final Ch1 and Ch2 datasets. Click-evoked ABRs included the same subjects and sound levels in both channels. Accuracy measures were computed for all sound levels tested, from 10-80 dB in 10 dB steps. All sound levels with fewer than 10 waveforms were excluded from training, analyses, and figures.

**Table 1:** Composition of the dataset by condition and time point. Table entries report the sex distribution, ketamine dosage, and the number of subjects contributing ABR data at each time point for the small-arms fire (SAF) and continuous noise exposure cohorts.

| Exposure Group Information |  |  |  |  |  |  |  |  |  |  |  |
| --- | --- | --- | --- | --- | --- | --- | --- | --- | --- | --- | --- |
|  |  | Sex |  | Ketamine Dosage |  | No. Subjects Included at Each Timepoint |  |  |  |  |  |
| SAF |  | M | F | 60 mg/kg | 40 mg/kg | Baseline | Day 2 | Day 7 | Day 14 | Day 28 | Day 56 |
|  | 80 dB | 7 | 12 | 19 | 0 | 19 | 17 | 18 | 14 | 10 | 5 |
|  | 131 dB | 1 | 1 | 2 | 0 | 2 | 0 | 2 | 2 | 2 | 2 |
|  | 134 dB | 15 | 16 | 27 | 4 | 31 | 17 | 27 | 21 | 16 | 17 |
|  | 137 dB | 5 | 6 | 11 | 0 | 11 | 11 | 11 | 8 | 2 | 2 |
|  | 140 dB | 7 | 3 | 10 | 0 | 10 | 6 | 8 | 3 | 4 | 4 |
| Continuous |  |  |  |  |  | Baseline | Week 1 | Week 2 | Week 3 | Week 4 |  |
|  | 87.5 dB | 4 | 4 | NA | NA | 8 | 8 | 7 | 0 | 3 |  |

**Table 2:** Composition of the training dataset by ABR stimulus frequency, stimulus level, and recording channel. Entries report the number of manually labeled waveforms used to train the Ch1 and Ch2 models.

| Dataset Composition |  |  |  |  |  |  |  |  |  |  |
| --- | --- | --- | --- | --- | --- | --- | --- | --- | --- | --- |
|  |  | 80 dB | 70 dB | 60 dB | 50 dB | 40 dB | 30 dB | 20 dB | 10 dB | Total |
| Channel 1 |  |  |  |  |  |  |  |  |  | 2863 |
|  | Click | 154 | 134 | 132 | 123 | 118 | 53 | 0 | 0 | 714 |
|  | 4 kHz | 128 | 115 | 132 | 98 | 43 | 0 | 0 | 0 | 516 |
|  | 8 kHz | 90 | 90 | 72 | 78 | 79 | 72 | 64 | 16 | 561 |
|  | 10 kHz | 73 | 77 | 69 | 92 | 69 | 67 | 63 | 14 | 524 |
|  | 16 kHz | 69 | 77 | 86 | 90 | 70 | 61 | 59 | 36 | 548 |
| Channel 2 |  |  |  |  |  |  |  |  |  | 2759 |
|  | Click | 154 | 134 | 132 | 123 | 118 | 53 | 0 | 0 | 714 |
|  | 4 kHz | 114 | 123 | 103 | 105 | 49 | 0 | 0 | 0 | 494 |
|  | 8 kHz | 75 | 73 | 82 | 72 | 75 | 60 | 57 | 17 | 511 |
|  | 10 kHz | 77 | 68 | 68 | 79 | 68 | 73 | 56 | 21 | 510 |
|  | 16 kHz | 62 | 83 | 72 | 69 | 73 | 78 | 62 | 31 | 530 |

Each response was treated as a discrete time sequence, with the indices of peaks and troughs annotated as the ground truth targets. For channel 1, waves I and III were labeled; for channel 2, waves I, IV, and V were labeled (Fig. ***1***B).

### CNN Architecture

Two separate CNNs were developed, with a dedicated network for each channel, both sharing a common architectural framework with minor structural variations. A detailed parameter comparison is summarized in Table 3. The input to each model was a two-channel tensor containing the primary and reference ABR signals. The primary signal was used for peak and trough prediction, while the other channel served as a contextual reference. Primary and reference roles were assigned according to the channel being predicted.

**Table 3:**
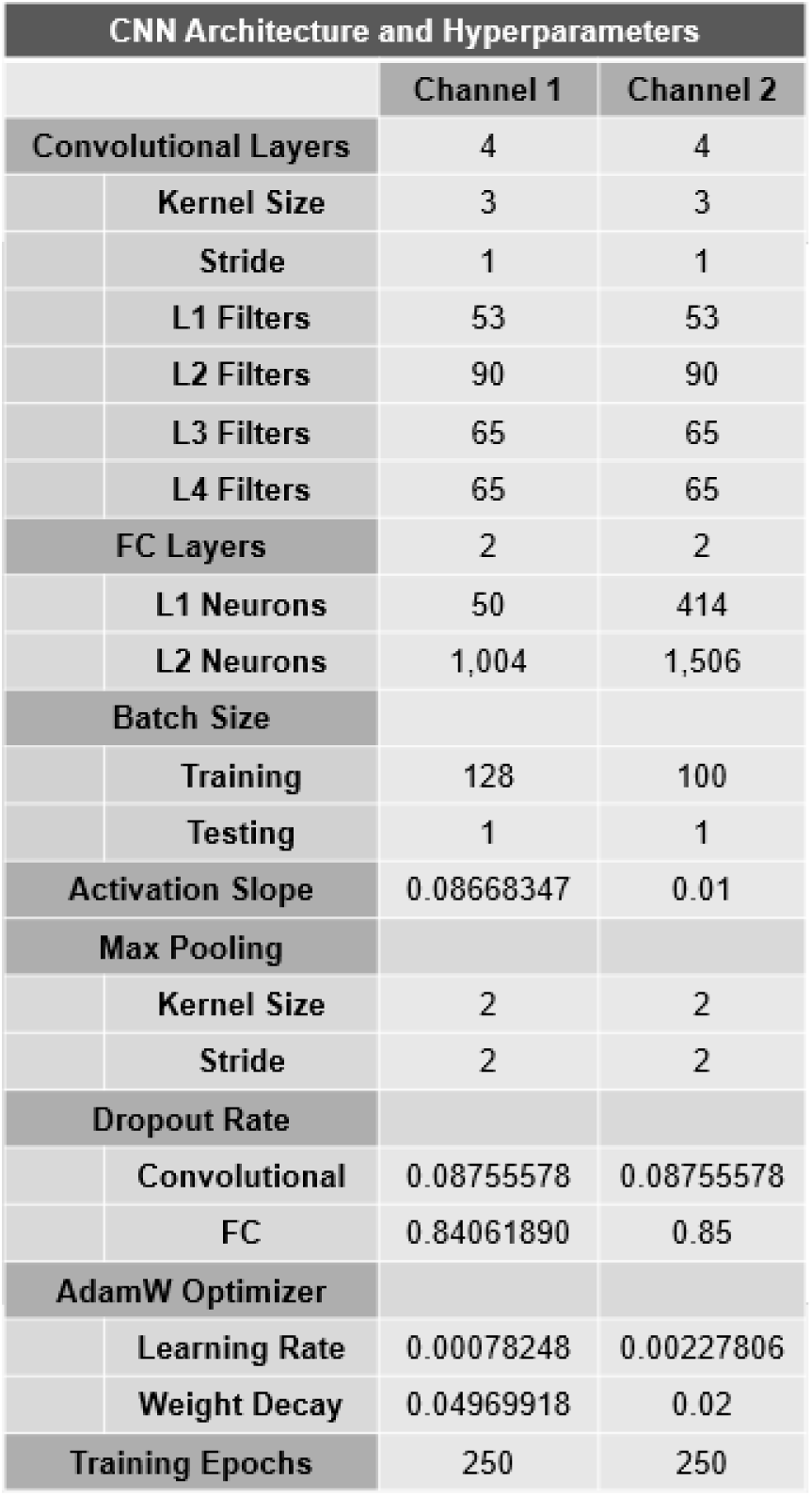
CNN architecture and hyperparameters for the Ch1 and Ch2 models. Entries report the final model configuration and hyperparameters used for training and inference.

Each network processed the input through four sequential convolutional blocks, followed by two feed-forward fully connected (FC) layers (Fig. 2). A block comprised a one-dimensional convolution, batch normalization, leaky rectified linear unit (LeakyReLU) activation, max pooling, and dropout. Batch normalization stabilized training by normalizing activations across the batch, while LeakyReLU introduced non-linearity to enable learning of complex patterns. Max pooling reduced temporal resolution while retaining dominant features and lowering computational load. Dropout was applied after every block to mitigate overfitting by randomly deactivating units during training. Relevant hyperparameters (dropout rate, activation slope, kernel stride, etc.) are shown in Table 3.

**Fig. 2:**
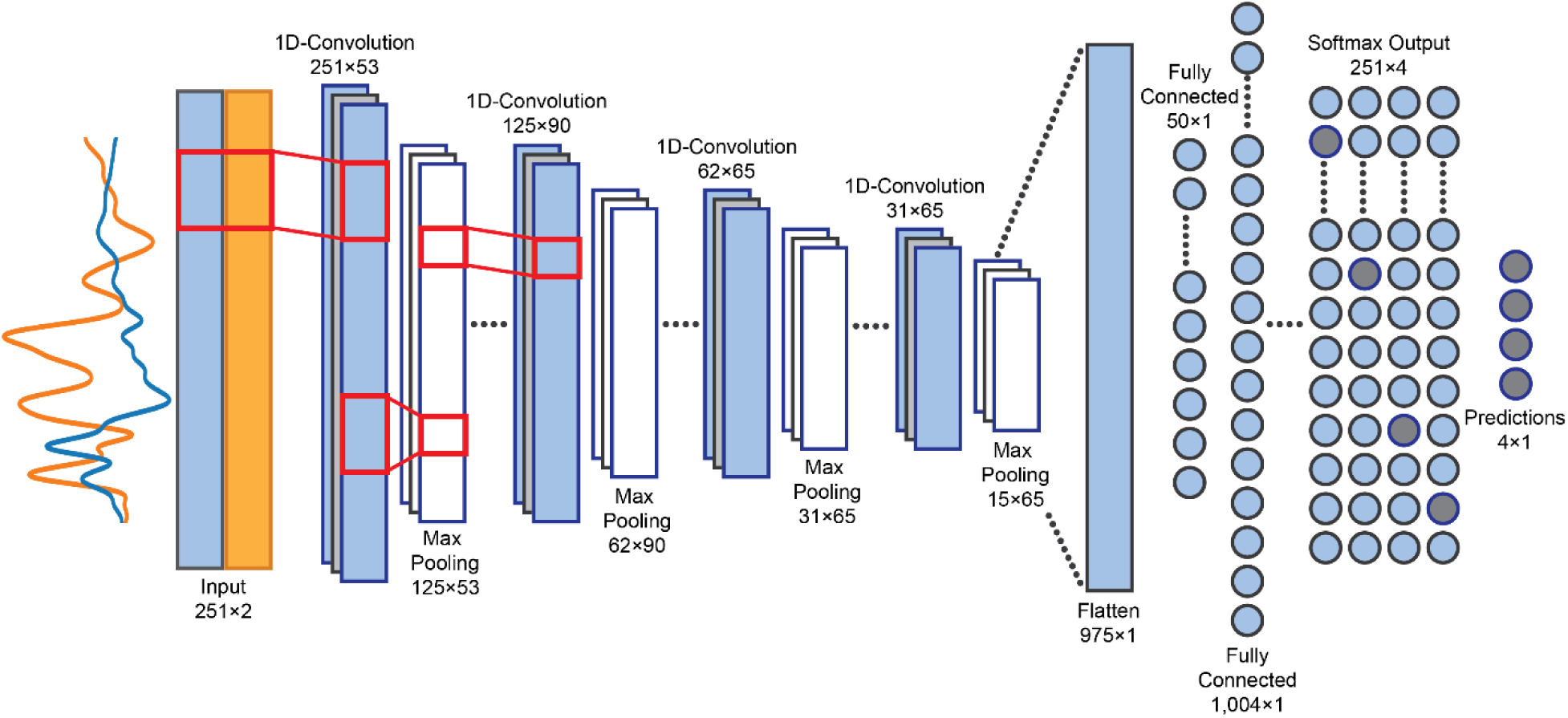
CNN architecture for the Ch1 model. The two-channel input (primary: Ch1, blue; reference: Ch2, orange) is jointly processed in the first convolutional layer, where each filter learns weighted combinations of both channels to produce one-dimensional feature maps. Each convolutional block comprises batch normalization, LeakyReLU activation, max-pooling for temporal down-sampling, and dropout for regularization/mitigating overfitting. The final feature maps are flattened and passed through fully connected layers with LeakyReLU, yielding nonlinear combinations of the learned features. The output layer generates softmax probability distributions for each target, with the index of maximum probability selected as the predicted location.

**Fig. 3:**
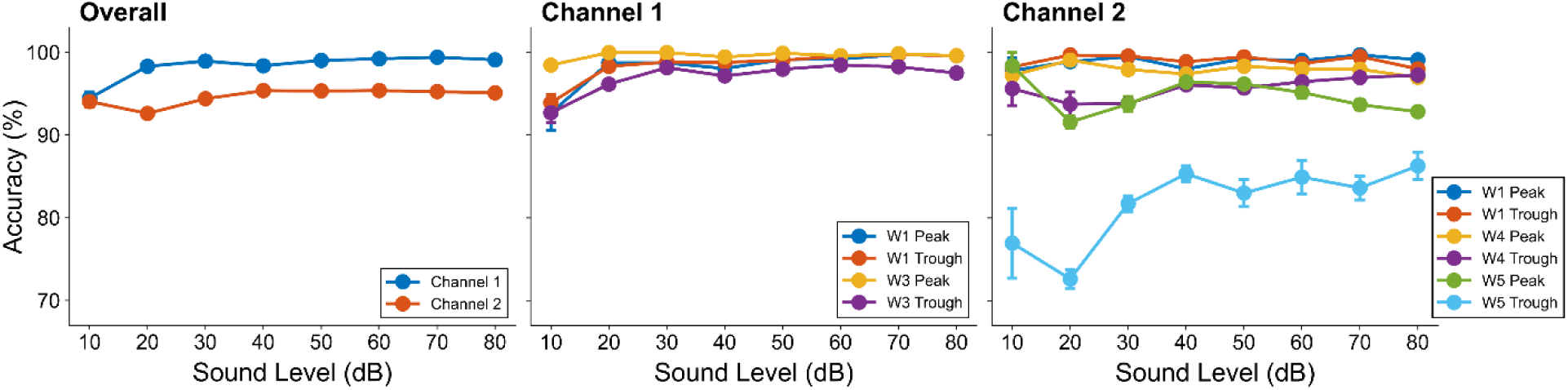
CNN prediction accuracy by stimulus level for all ABR stimuli. Mean accuracies were calculated from pooled test data from 10-fold cross-validation (CV) for Ch1 and Ch2 and further stratified by waveform target for each channel. Overall accuracy was 98.84% for Ch1 and 95.50% for Ch2, with variation ≤0.7% across runs. Accuracy remained stable across sound levels for both channels; however, target-level analysis indicated that the lower accuracy in Ch2 was primarily attributable to the wave V trough.

### Model Output and Prediction Strategy

Following the convolutional stack, the resulting time-feature representation was flattened (reshaped into a single vector) and passed through FC layers with LeakyReLU activations, yielding nonlinear combinations of features across time points and feature maps. For each target, the network produced a logit vector over discretized time indices (four targets for Ch1; six for Ch2, corresponding to a peak and trough for each wave of interest). These logits were unnormalized scores representing the relative evidence for an event at each index. A per-target softmax function, applied along the time dimension, converted logits to probabilities, and the index with the highest probability was taken as the initial prediction. No ties occurred in this dataset; if present, ties would default to the earlier index.

To further refine these predictions, a channel-specific post-processing target adjustment (TA) was applied. The TA function first verified whether the predicted index corresponded to a local extremum (maximum for peaks, minimum for troughs). If not, a ±0.82 ms window (±20 indices) around the initial prediction was searched for the nearest valid extremum, except for channel 2 peaks, where the prediction was instead moved to the most prominent maximum within the window.

To preserve the temporal order of the targets, the eligible range within each search window was further restricted. For channel 1, the wave III peak was adjusted first and used as an anchor: the wave III trough search was limited to indices after the adjusted wave III peak, the wave I peak search to indices before it, and the wave I trough search to indices between the adjusted wave I and wave III peaks. For channel 2, targets were adjusted sequentially from wave I through wave V, with the lower limit of each search window set immediately after the preceding adjusted target. If no extremum was found within the eligible range, the original prediction was retained. By enforcing alignment with true extrema, the function ensured precise peak and trough localization for subsequent ABR analyses.

### Training Procedure

Training used 10-fold cross-validation (CV). Each iteration used 90% of the data for training and 10% for testing, with mutually exclusive test sets. Folds 1-9 contained 289 signals for Ch1 and 284 for Ch2, while fold 10 contained 295 signals for Ch1 and 280 for Ch2. Data was randomly assigned to these splits for each training run.

Models were trained for 250 epochs, minimizing cross-entropy loss on the initial (pre-adjustment) predictions. Training batch size was 128 for Ch1 and 100 for Ch2, using AdamW optimization with decoupled weight decay. The base learning rate was not scheduled (learning rate and weight decay are reported in Table 3). During evaluation, a batch size of 1 was used to enable per-sample diagnostics. Evaluation was performed with dropout disabled and batch normalization using fixed running statistics.

### Evaluation Metrics

Overall CNN performance was evaluated based on post-adjustment prediction accuracy. A prediction was considered correct only if the adjusted index exactly matched the ground truth annotation index. Outcomes were aggregated across folds, and global accuracy metrics were computed after CV completion. CV ensured that all data contributed to both training and evaluation, reducing sensitivity to any single fold containing noisy or ambiguous signals.

### Hyperparameter Optimization

Initial hyperparameters were selected with the Hyperband algorithm, an adaptive resource-allocation (multi-armed bandit) method that trains many configurations with small budgets and progressively allocates more resources to promising candidates, minimizing validation loss (Li et al. 2018). Hyperparameter optimization was implemented using the Optuna Python Library (Akiba et al. 2019). After automated tuning, limited manual refinement was performed based on validation performance. Final hyperparameters are listed in Table 3.

### Dataset Ablation

To assess sensitivity to training-set size, dataset ablation was performed by systematically reducing the number of ABR signals used for model training. Analysis was conducted separately for the two channel models. Removal levels ranged from 0% to 95% in 5% increments, with 10 independent repeats per condition. For a given level, the same fraction of signals was excluded from every stimulus group. For each reduced dataset, the CNN was trained and evaluated using the 10-fold cross-validation procedure and hyperparameters described above. Performance was summarized by waveform target using mean absolute error for initial and adjusted predictions, as well as exact accuracy after target adjustment. Mean absolute error was calculated as the absolute distance between predicted and ground-truth indices. Metrics were averaged across repeats, with variability reported as the standard deviation.

### Statistical Analysis

Statistical analyses were conducted in MATLAB (MathWorks; R2024b). To evaluate CNN robustness across ABRs associated with distinct auditory damage profiles, prediction accuracy in the SAF cohort was analyzed using binomial generalized linear mixed-effects regression with a logit link (fitglme). Each signal was scored as the number of correctly predicted targets out of the total number of evaluated targets.

Fixed model effects were damage condition (functional versus sham) and stimulus sound level as categorical fixed effects, with subject included as a random intercept to account for repeated observations within animals. A secondary model additionally included stimulus frequency and exposure day as categorical fixed effects. Models were fit using the Laplace approximation. Statistical significance of the group effect was assessed using two-sided Wald tests at α = 0.05. Likelihood-ratio tests comparing models with and without the group term were also reported. Adjusted odds ratios (OR) and 95% confidence intervals (CI) were calculated from the group coefficient. Observed accuracy was calculated as the total number of correctly predicted targets divided by the total number evaluated within each group. The observed group difference was reported in percentage points as sham accuracy minus functional accuracy.

### Hardware and Software Environment

Model development and training were conducted on a Dell Precision 5560 laptop equipped with an 11th Gen Intel Core i7-11800H CPU, an NVIDIA T1200 Laptop GPU, and 16 GB of RAM. The system ran Windows 10 Pro (version 10.0.19045) with 474 GB of available storage. CUDA version 12.7 was used for GPU acceleration. All CNN code was implemented in Python (version 3.12.9) using the PyTorch deep learning framework (version 2.6.0).

## Results

Manually labeled peaks from channel 1 (waves I, III) and channel 2 (waves I, IV, V) were used to train a CNN to predict discrete time indices for each waveform feature. The training data included both SAF-exposed animals (damaged and sham) and continuous noise-exposed animals to increase robustness to hearing pathology.

### Accuracy

Aggregated across all ABR stimuli and labeled targets, accuracy over the 10 repeated CV runs was 98.90% (±0.04) for channel 1 and 95.00% (±0.25) for channel 2. Results were stable across stimulus levels and absolute ABR amplitudes in both channels (3, left). For individual targets, channel 1 remained consistent across waves and sound levels (3, middle); channel 2 accuracy was also quite consistent except for some misidentification of the wave V trough, which accounted for most errors (3, right).

Stimulus-specific accuracy was similarly consistent, with high performance across click and pure-tone conditions (Fig. 4). Values ranged from 98.16% (± 0.22) to 99.52% (±0.09) for channel 1 and 93.47% (±0.24) to 97.01% (±0.17) for channel 2 (click: 99.00% (±0.05)/ 97.01% (±0.17); 4 kHz: 98.16% (±0.22)/94.12% (±0.48); 8 kHz: 98.40% (±0.07)/ 94.06% (±0.31); 10 kHz: 99.52% (±0.09)/95.58% (±0.52); 16 kHz: 99.37% (±0.06)/93.47% (±0.24); Ch1/Ch2). Residual errors were concentrated at the wave V peak and trough, with a localized reduction for the 4 kHz stimulus at 40 dB, which is likely due to this being near threshold for 4 kHz.

**Fig. 4:**
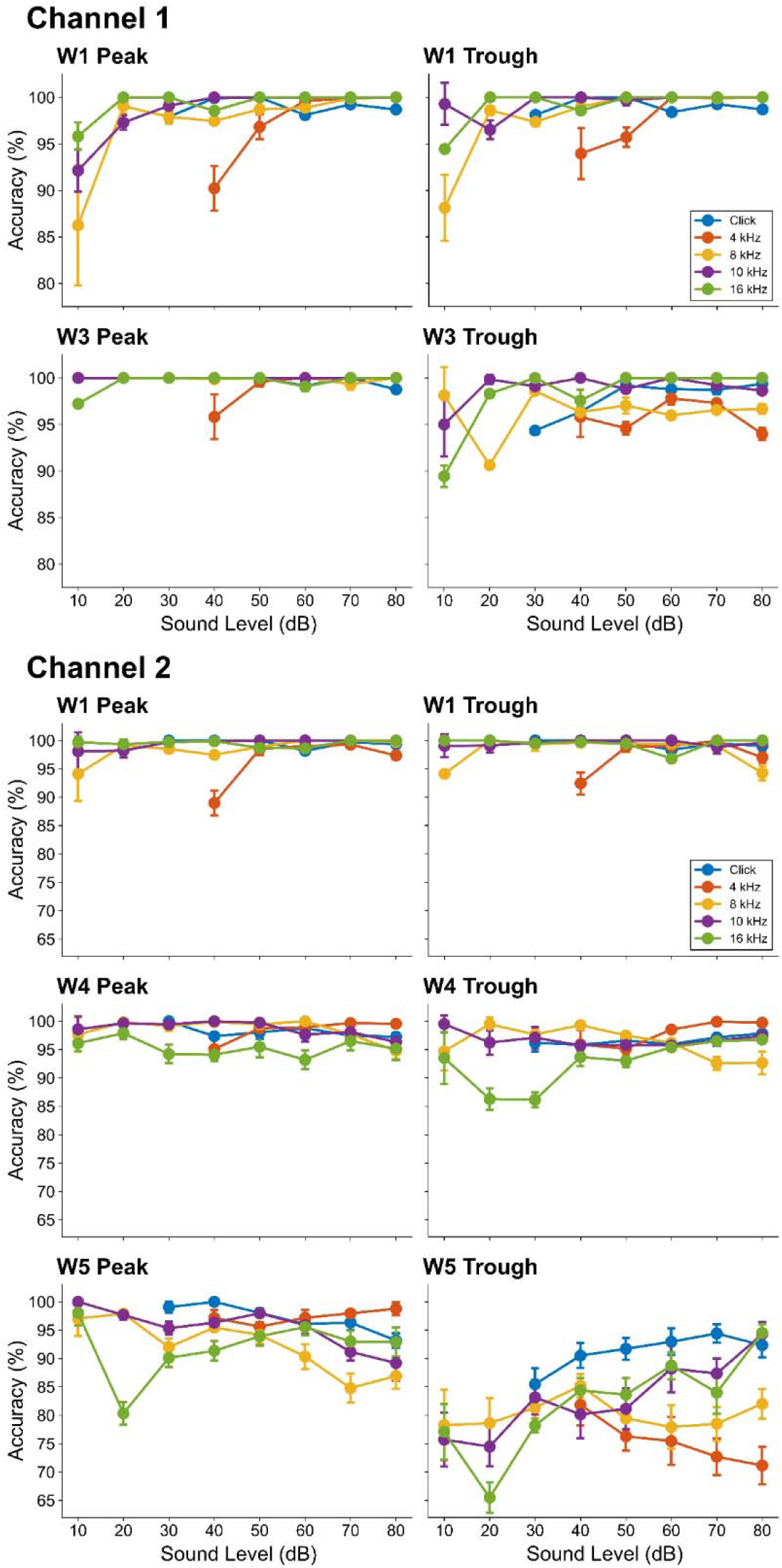
Stimulus-specific prediction accuracy for each waveform target in Ch1 and Ch2. Consistent with the overall trends, accuracy was generally stable across stimuli and sound levels, with reduced performance concentrated at the wave V peak and trough and near threshold sound levels.

Prediction accuracy remained high in both functionally impaired (threshold shift) and non-impaired (no threshold shift) groups (Fig. 5), although group effects differed by channel. Subjects underwent high-intensity SAF noise exposure (sham: 80 dB pSPL; experimental: 131-140 dB pSPL) and were classified as functional if they exhibited a ≥10 dB threshold shift 7 days post-insult; baseline ABRs were included for each subject.

**Fig. 5:**
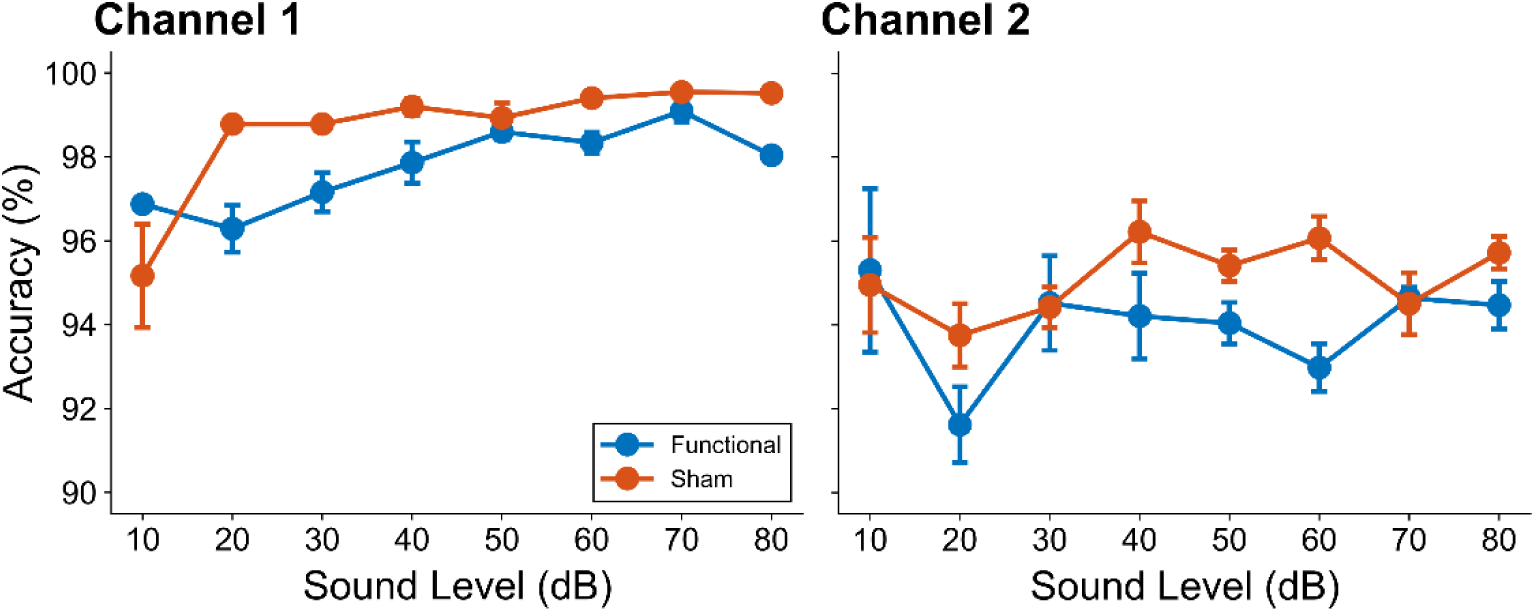
Prediction accuracy for small-arms fire (SAF) noise cohorts. Subjects underwent high-intensity noise exposure (sham: 80 dB SPL; experimental: 131-140 dB SPL) and were classified as functional if they exhibited a ≥10 dB threshold shift 7 days post-exposure. Accuracy remained high in both groups, but sham animals showed significantly higher prediction accuracy than functional animals in channel 1 (P = 0.0313; observed: 99.31% vs. 98.31%; model-estimated mean difference: 0.81 percentage points) and channel 2 (P = 0.0443; observed: 95.86% vs. 94.44%; model-estimated mean difference: 1.29 percentage points). This effect remained significant in a secondary model additionally adjusted for stimulus frequency and exposure day in channel 1 (P = 0.0077) and channel 2 (P = 0.0276).

**Fig. 6:**
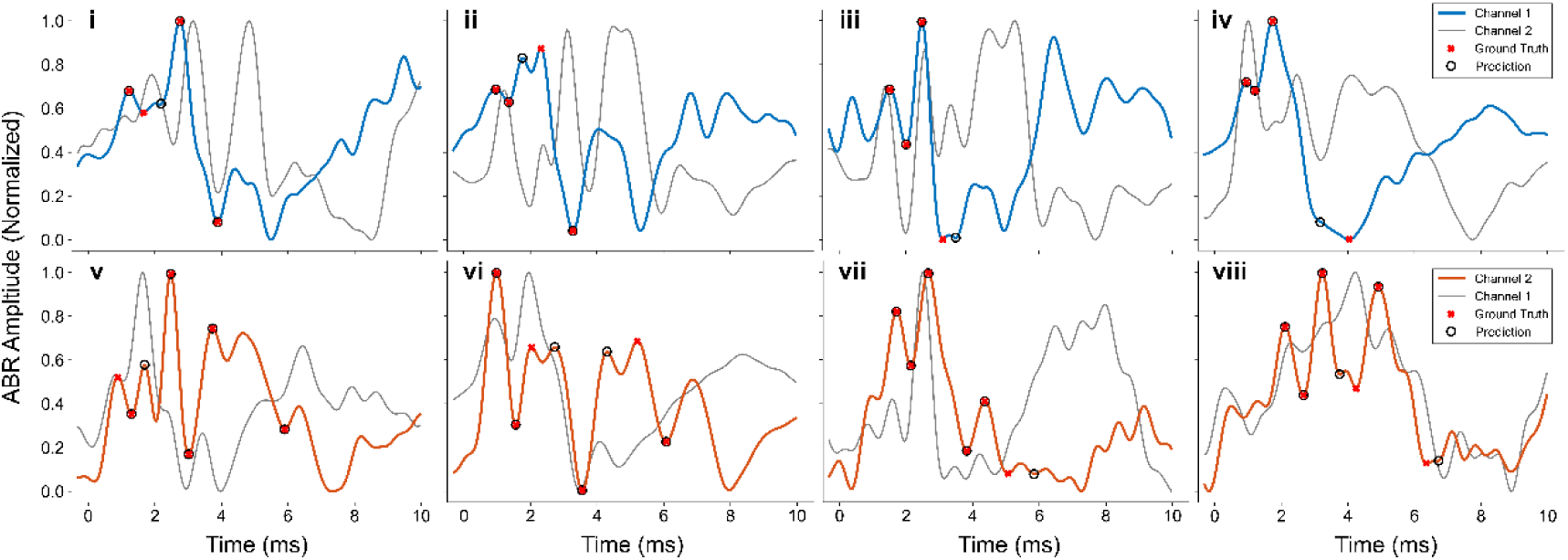
Examples of incorrect predictions for Ch1 and Ch2 (i: 4 kHz, 50 dB; ii: click, 80 dB; iii: 8 kHz, 70 dB; iv: 8 kHz, 70 dB; v: 4 kHz, 70 dB; vi: 16 kHz, 70 dB; vii: 8 kHz, 30 dB; viii: 8 kHz, 10 dB). Split peaks (ii, wave III; v, waves I and V; vi, waves IV and V) were associated with increased error likelihood and were more prevalent in Ch2, contributing to its lower overall accuracy. The Ch2 wave V trough was the most frequent error, consistent with its variable morphology across subjects and stimulus levels. At higher stimulus levels (v, vi), a distinct wave V knee may be labeled as the trough, whereas at lower levels (vii, viii) this feature becomes less distinct and merges int noise, complicating both manual labeling and CNN prediction.

In channel 1, binomial generalized linear mixed-effects regression (see Methods) indicated higher accuracy in sham than in functional animals (Wald P = 0.0348; likelihood-ratio P = 0.0399). The observed difference was small (99.08% versus 98.23%), although the adjusted odds of a correct prediction were higher in sham animals (OR, 3.36; 95% CI, 1.09-10.33). Channel 2 showed the same direction of effect but was not significant in the primary model (95.25% versus 94.02%; OR, 1.24; 95% CI, 0.96-1.60; Wald P = 0.1058; likelihood-ratio P = 0.1123). In the secondary model additionally adjusted for stimulus frequency and exposure day, the group effect was significant by both tests in channel 1 (OR, 5.43; 95% CI, 1.77-16.64; Wald P = 0.0031; likelihood-ratio P = 0.0051) and channel 2 was on the edge of significance for a 95% null rejection threshold (OR, 1.28; 95% CI, 1.00-1.64; Wald P = 0.0470; likelihood-ratio P = 0.0530).

To assess applicability to conventional single-channel ABR recordings, a single-channel input CNN was also evaluated using only the primary signal (without the contextual reference channel). Performance remained similar to the two-channel model, with overall accuracies of 98.84% (±0.03) for channel 1 (standard rodent ABR configuration) and 94.28% (±0.30) for channel 2 (click: 98.90% (±0.08)/96.45% (±0.36); 4 kHz: 97.93% (±0.12)/93.14% (±0.42); 8 kHz: 98.41% (±0.09)/93.38% (±0.35); 10 kHz: 99.48% (±0.07)/94.93% (±0.38); 16 kHz: 99.44% (±0.07)/92.69% (±0.35); Ch1/Ch2).

### Incorrect Examples

Although errors were uncommon, channel 1 most frequently misclassified the wave III trough, which corresponds to just after the channel 2 wave IV peak (Parthasarathy and Bartlett 2012). These errors typically arose from ambiguous morphology, including closely spaced local minima and delayed, ill-defined minima (6iii, iv). Split peaks (double maxima separated by a shallow notch) were another prominent source of misidentification for both channels (6ii: wave III; 6v: wave V; 6vi: waves IV, V), particularly when adjacent peaks had similar amplitudes. Channel 2 had a higher incidence of split peaks, contributing to lower accuracy.

The most difficult target was the wave V trough, whose morphology varied substantially across stimulus levels and subjects, complicating manual labeling and increasing prediction variability. At higher levels, a distinct wave V knee may be labeled as the trough, whereas at lower levels this feature becomes less distinct and merges into noise (6v-viii). Consequently, even carefully curated evaluation sets can include cases where the trough location is inherently ambiguous, limiting annotation consistency.

### Target time adjustment

A post-processing target-adjustment (TA) function was applied as described in Methods. Because predictions were defined on discrete sample indices, shifts could occur only in integer steps (1 index ≈ 0.0410 ms, or 1/24414.0625 s). Among correct predictions (post-adjustment index = ground truth), TA distances were calculated separately for each CV run and summarized across runs. Mean of run-median adjustment distances ranged from 0-0.0819 ms across waveform targets and 0-0.0410 ms across stimuli, indicating minimal temporal correction overall (Table 4). Means of run-mean adjustments, stratified by waveform target, were likewise small, except for elevated values at the channel 1 wave III trough (0.0889 ms) and the channel 2 wave V trough (0.0852 ms) (Table 4; Fig. 7A). By stimulus, shifts were similarly stable, with a modest increase at 4 and 8 kHz in channel 1 (0.0652 and 0.0569 ms, respectively) (Table 4; Fig. 7B).

**Fig. 7:**
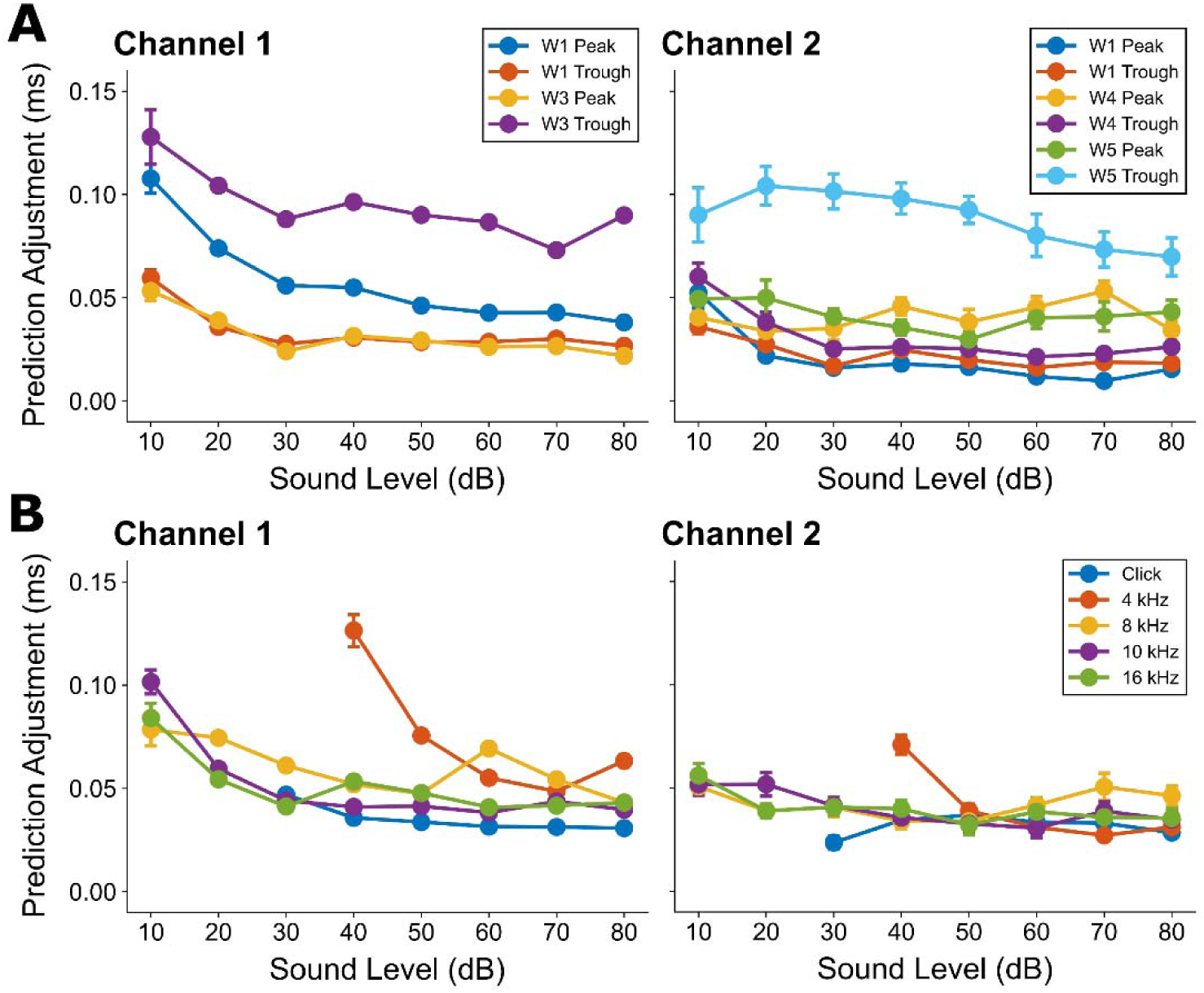
Average absolute adjustment distance applied by the target-adjustment (TA) function for ABR predictions. Means were calculated from pooled test data across 10-fold cross-validation (CV) and included only correct predictions (post-adjustment index = ground truth). The TA function checks whether the initial prediction lies on a local extremum (maximum for peaks, minimum for troughs); otherwise, it searches within ±0.82 ms (0.041 ms/index) and snaps to the nearest extremum. For Ch2 peaks only, the TA instead snaps to the largest maximum within this window. (A) Values for all ABR stimuli, stratified by waveform target; (B) values for all waveform targets, stratified by ABR stimulus, both shown as a function of sound level. Distances were generally stable and rose slightly at the lowest sound levels or near the 4 kHz threshold.

**Table 4:** Descriptive statistics for target-adjustment (TA) distances for correct predictions (post-adjustment index = ground truth). The top section is stratified by waveform target (all ABR stimuli combined), and the bottom section is stratified by stimulus (all waveform targets combined). As predictions are defined on discrete sample indices, TA shifts can occur only in integer index steps (1 index is approximately 0.0410 ms); therefore, observed distances indicate minimal temporal correction.

| Target Adjustment Distance – Correct Predictions Only – All Stimuli |  |  |  |  |  |  |
| --- | --- | --- | --- | --- | --- | --- |
|  |  | Mean of Run Means (ms) | Mean of Run Medians (ms) | SD Across Runs | Mean Within-Run SD | Median Sample Size per Run |
| Channel 1 |  |  |  |  |  |  |
|  | W1 Peak | 0.0487 | 0.0410 | 0.0005 | 0.0609 | 2834 |
|  | W1 Trough | 0.0299 | 0.0369 | 0.0008 | 0.0494 | 2839 |
|  | W3 Peak | 0.0280 | 0.0164 | 0.0003 | 0.0438 | 2855 |
|  | W3 Trough | 0.0889 | 0.0819 | 0.0012 | 0.0953 | 2797 |
| Channel 2 |  |  |  |  |  |  |
|  | W1 Peak | 0.0158 | 0.0000 | 0.0011 | 0.0463 | 2734 |
|  | W1 Trough | 0.0201 | 0.0000 | 0.0007 | 0.0493 | 2732 |
|  | W4 Peak | 0.0419 | 0.0000 | 0.0027 | 0.1198 | 2702 |
|  | W4 Trough | 0.0262 | 0.0000 | 0.0011 | 0.0570 | 2653 |
|  | W5 Peak | 0.0393 | 0.0041 | 0.0037 | 0.1022 | 2608 |
|  | W5 Trough | 0.0852 | 0.0410 | 0.0064 | 0.1359 | 2297 |

| Target Adjustment Distance – Correct Predictions Only – All Targets |  |  |  |  |  |  |
| --- | --- | --- | --- | --- | --- | --- |
|  |  | Mean of Run Means (ms) | Mean of Run Medians (ms) | SD Across Runs | Mean Within-Run SD | Median Sample Size per Run |
| Channel 1 |  |  |  |  |  |  |
|  | Click | 0.0335 | 0.0410 | 0.0004 | 0.0510 | 2827.5 |
|  | 4 kHz | 0.0652 | 0.0410 | 0.0013 | 0.0891 | 2026 |
|  | 8 kHz | 0.0569 | 0.0410 | 0.0008 | 0.0768 | 2208 |
|  | 10 kHz | 0.0451 | 0.0410 | 0.0011 | 0.0621 | 2086 |
|  | 16 kHz | 0.0482 | 0.0410 | 0.0006 | 0.0649 | 2178 |
| Channel 2 |  |  |  |  |  |  |
|  | Click | 0.0324 | 0.0000 | 0.0019 | 0.0919 | 4153.5 |
|  | 4 kHz | 0.0357 | 0.0000 | 0.0023 | 0.0823 | 2791 |
|  | 8 kHz | 0.0413 | 0.0000 | 0.0025 | 0.1039 | 2888.5 |
|  | 10 kHz | 0.0381 | 0.0000 | 0.0028 | 0.0909 | 2923 |
|  | 16 kHz | 0.0386 | 0.0000 | 0.0023 | 0.0967 | 2972.5 |

Empirical cumulative distribution functions (ECDFs) confirmed that TA distances were strongly concentrated near zero in both channels (Fig. 8). The 90th percentile adjustment was calculated separately for each CV run and summarized as the mean ± standard deviation across runs (Table 5). For most targets, the 90th percentile was <0.0819 ms, indicating that at least 90% of correct predictions required index adjustments ≤ 2. Larger cutoffs were limited to the channel 1 wave I peak and wave III trough, and most prominently to the channel 2 wave V trough. For the small proportion of incorrect predictions (post-adjustment index ≠ ground truth), absolute deviations were generally 0.7-1.05 ms (Table 5). Here, the CNN was basically guessing, with approximately 40% of adjustments going farther away from the manually identified peak (not shown).

**Fig. 8:**
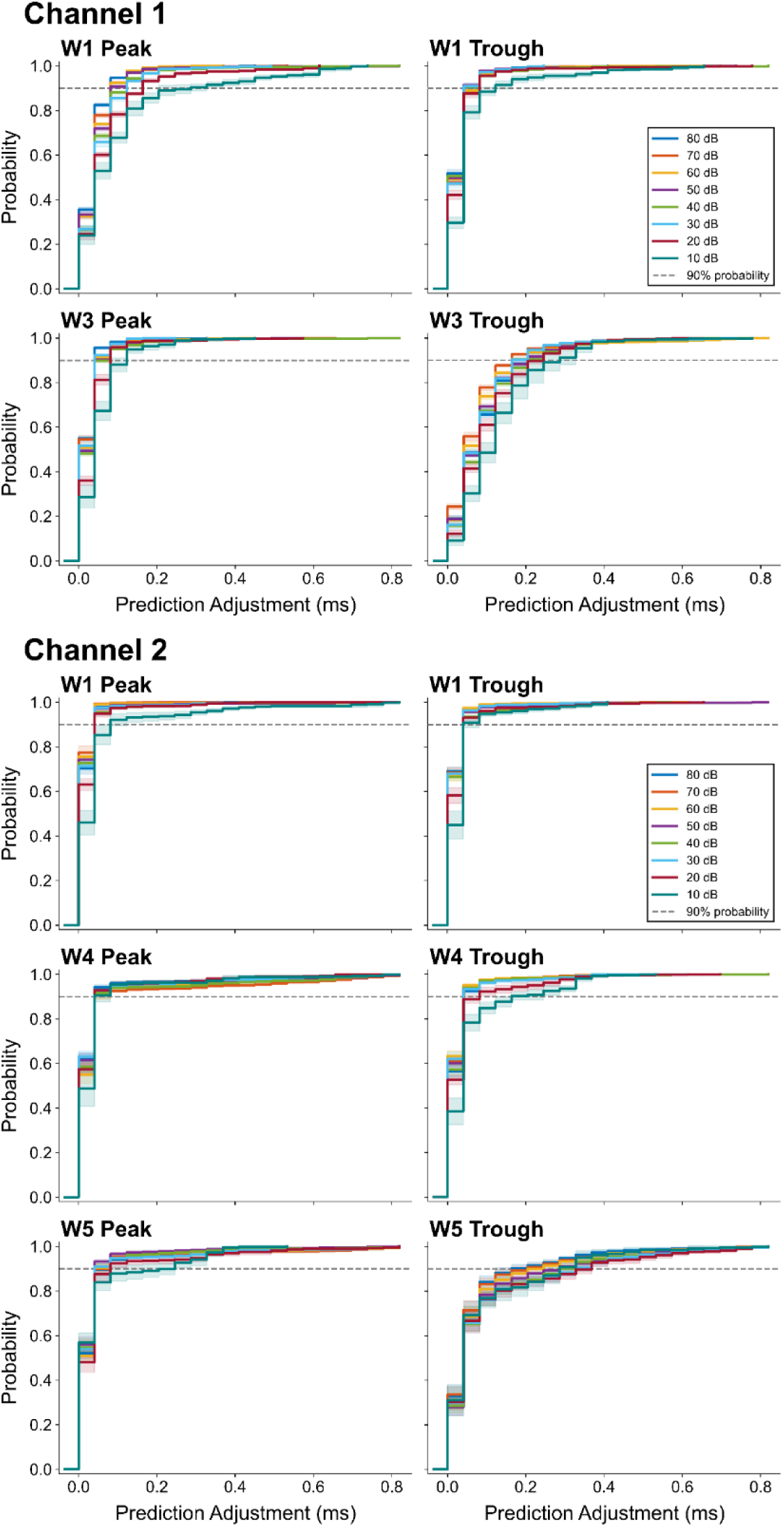
Empirical cumulative distribution functions (ECDFs) of adjustment distances applied by the target-adjustment (TA) function for Ch1 and Ch2 across all ABR stimuli. ECDFs were computed from pooled test data across 10-fold cross-validation (CV) and restricted to correct predictions (post-adjustment index = ground truth). Distances were sorted, and cumulative probabilities were incremented by 1/n per observation (n = total samples). The gray line denotes the 90th percentile adjustment distance.

**Table 5:** Descriptive statistics of post-adjustment prediction error for incorrect predictions (post-adjustment index ≠ ground truth) across all ABR stimuli, reported by waveform target. Distances reflect the absolute error between the target-adjusted prediction and the ground-truth index.

| Distance from Ground Truth – Incorrect Predictions Only – All Stimuli |  |  |  |  |  |  |
| --- | --- | --- | --- | --- | --- | --- |
|  |  | Mean of Run Means (ms) | Mean of Run Medians (ms) | SD Across Runs | Mean Within-Run SD | Median Sample Size per Run |
| Channel 1 |  |  |  |  |  |  |
|  | W1 Peak | 0.8900 | 0.8602 | 0.0299 | 0.2954 | 29 |
|  | W1 Trough | 0.7908 | 0.7434 | 0.0273 | 0.3399 | 24 |
|  | W3 Peak | 0.7777 | 0.7598 | 0.0539 | 0.2246 | 8 |
|  | W3 Trough | 0.7571 | 0.7086 | 0.0145 | 0.2657 | 66 |
| Channel 2 |  |  |  |  |  |  |
|  | W1 Peak | 0.9748 | 0.8233 | 0.0344 | 0.3476 | 25 |
|  | W1 Trough | 0.8146 | 0.7352 | 0.0525 | 0.4563 | 27 |
|  | W4 Peak | 0.9867 | 0.9134 | 0.0226 | 0.2709 | 57 |
|  | W4 Trough | 0.8372 | 0.8110 | 0.0189 | 0.3200 | 106 |
|  | W5 Peak | 1.0542 | 1.0199 | 0.0137 | 0.2407 | 151 |
|  | W5 Trough | 1.4338 | 1.3640 | 0.0167 | 0.5998 | 462 |

**Table 6:**
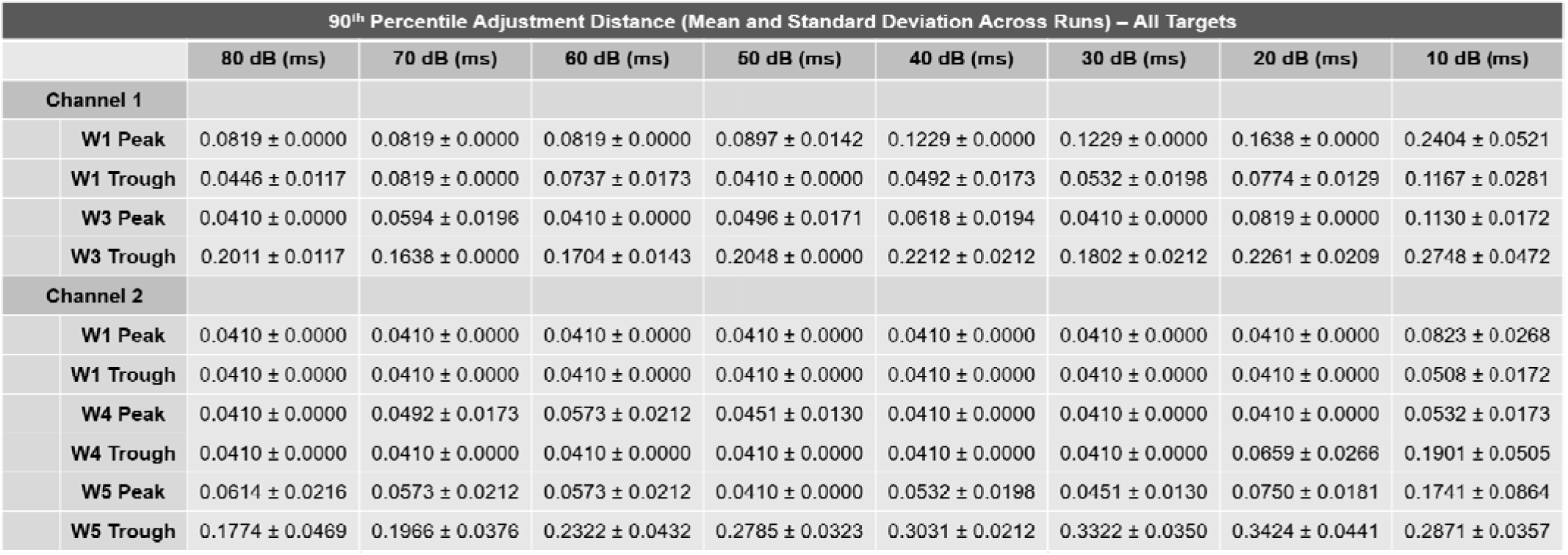
90th percentile of absolute target-adjustment (TA) distance for correct predictions, stratified by waveform target and stimulus sound level. Values are reported in milliseconds and were calculated from predictions for which the post-adjustment index matched the ground-truth index. Because predictions were defined on discrete sample indices, adjustment distances occurred in increments of approximately 0.0410 ms.

### Calibration

Calibration curves were constructed from the initial (non-adjusted) classification outputs for each target in both channel models (Fig. 9) to assess the relationship between confidence and accuracy. Softmax scores at the predicted indices and corresponding correctness labels were extracted from test data, and mean predicted probabilities were plotted against empirical accuracies using five quantile-based bins with equal sample counts. The diagonal represents perfect calibration, where predicted confidence matches observed accuracy. Both models were generally well calibrated: channel 2 closely followed the diagonal across targets, with a slight downward deviation for the wave V trough, whereas channel 1 curves tended to lie slightly above the diagonal, consistent with mild underconfidence.

**Fig. 9:**
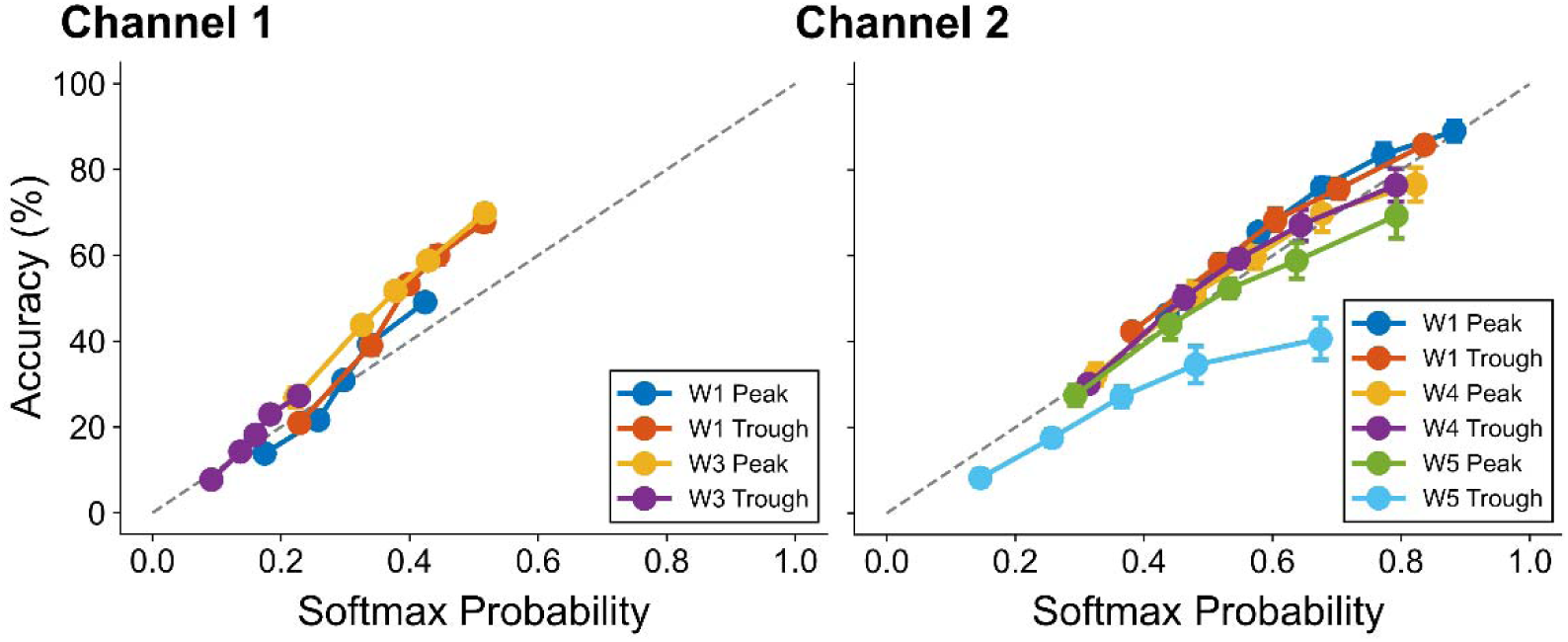
Calibration curves for initial (non-adjusted) predictions of each waveform target in the Ch1 and Ch2 models, for all ABR stimuli. Softmax confidence scores and correctness labels were extracted from test outputs. Mean predicted probabilities were plotted against empirical accuracies in five quantile-based bins (equal sample counts). The diagonal indicates perfect calibration. Both models were generally well calibrated: Ch2 closely followed the diagonal across targets, with a slight downward deviation for the wave V trough, while Ch1 curves were slightly above the diagonal, consistent with mild underconfidence.

### Dataset Ablation

Dataset ablation was used to evaluate sensitivity to training-set size (Fig. 10). Initial and target-adjusted mean absolute error remained low across removal levels, and post-adjustment accuracy was largely preserved. Performance declined only after substantial data reduction, beginning at ∼90% removal for all targets except the wave V trough (channel 1, 289 remaining signals, and channel 2, 280 remaining signals). Overall, the CNNs maintained reliable ABR labeling despite significant reductions in training data.

**Fig. 10:**
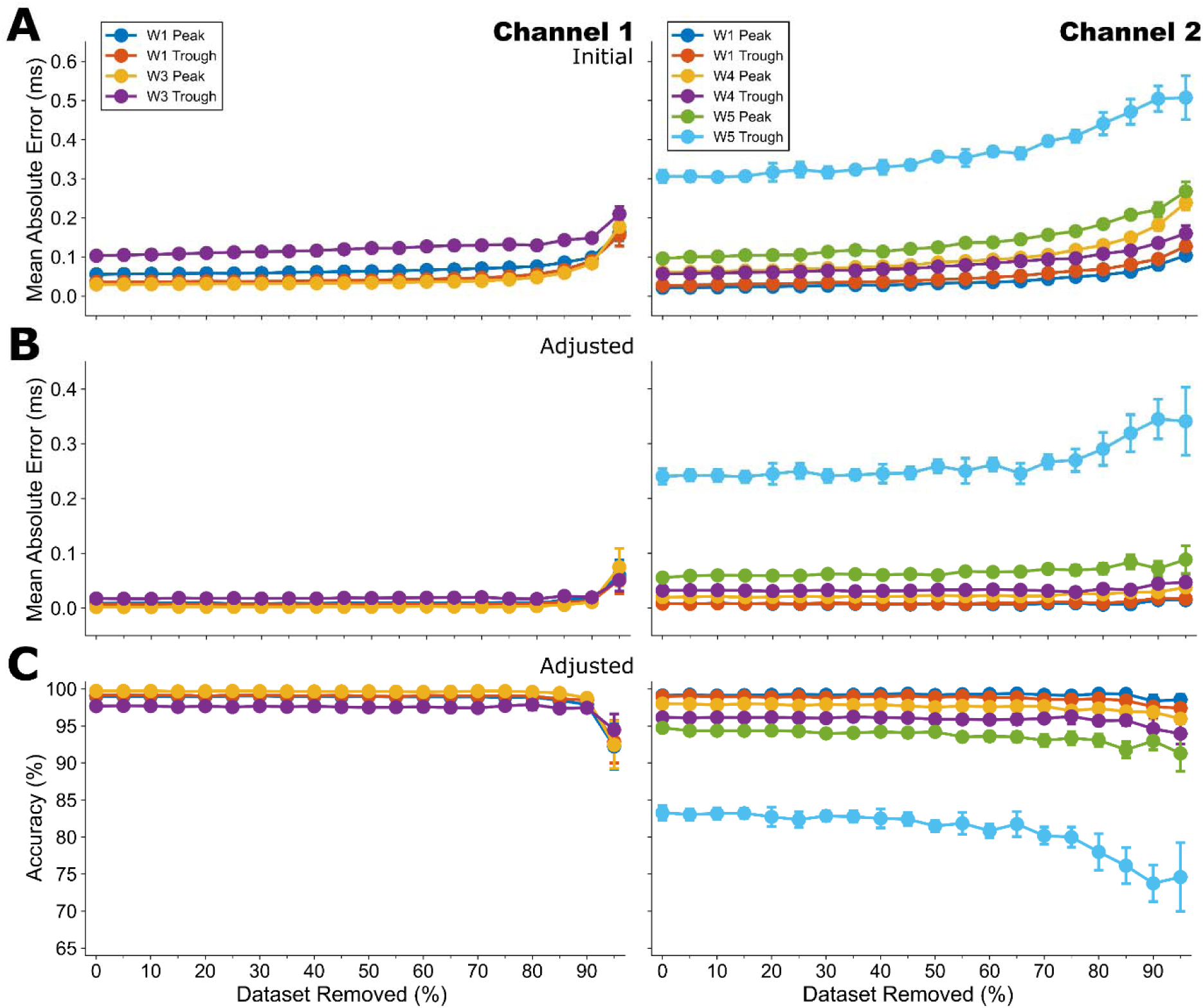
Dataset ablation analysis of CNN prediction performance. Model robustness was evaluated using stimulus-stratified ablation, with progressively larger fractions of ABR signals removed evenly across frequency groups. Mean absolute error (average absolute distance between predicted and ground-truth location) is shown for the initial CNN outputs (A) and target-adjusted outputs (B), with adjusted accuracy shown in C. Performance remained stable across most ablation levels and declined only after ∼90% data removal (Ch1: 90% removal, 289 remaining signals; Ch2: 90% removal, 284 remaining signals).

### Computing Timing

Timing metrics were obtained using the full labeled dataset for each channel model (Ch1: n=2,863; Ch2: n=2,759). Cross-validation runtime was defined as the total time required to train all 10 folds using the final hyperparameter configuration (Ch1: 7 m 6.51 s; Ch2: 8 m 36.30 s). Final model training time was defined as the time required to train each model once on its full dataset (Ch1: 45.74 s; Ch2: 56.64 s). After training, the saved model state was reloaded in evaluation mode (dropout disabled and batch normalization fixed) for inference benchmarking. Per-signal inference time was measured at batch size 1 using 600 randomly sampled signals, with 100 used for warm-up and 500 timed individually; mean inference time was 1.08384 ± 0.111 ms/signal for channel 1 and 1.2120 ± 0.0572 ms/signal for channel 2.

## Discussion

### Summary of results

A CNN-based classifier was developed to automate the identification of two-channel ABR peaks and troughs in a rodent model that included normal hearing and noise-exposed animals with threshold shifts. Accuracy was consistently high across stimulus types, sound levels, and auditory injury profiles (Fig. 3), demonstrating robustness to waveform heterogeneity. Residual errors were concentrated within the channel 2 wave V trough, which accounted for a disproportionate share of misclassifications, and to a lesser degree in the wave V peak (Fig. 4). Within the SAF-noise-exposed cohort, performance remained strong in both functionally impaired animals with threshold shifts and non-impaired animals. Although the functional group showed a statistically significant reduction in accuracy relative to the sham group in both channels, the magnitude of the disparity was small, suggesting that auditory impairment modestly increased peak-picking difficulty without substantially reducing overall accuracy. Comparable accuracy in the single-channel input model suggests that the classifier derived most of its predictive information from the primary signal alone, supporting its use in conventional single-channel ABR configurations. Training of the CNN only required a few hundred identified peaks to achieve good accuracy (Fig. 10). Once trained, inference required approximately one ms of computational time per signal using personal computers, making the algorithm well-suited to large datasets and longitudinal studies in which manual labeling would be prohibitively time-consuming.

### Comparison with previous studies

Automated ABR evaluation has been a goal in clinical and research auditory research for decades (e.g. Gronfors 1993). Many studies have used automated ABR detection for the creation of an audiogram across age (Liu et al. 2025, Zhang et al. 2026, Wang et al. 2021, McKearney et al. 2022, Ma et al. 2023), and although clinically quite useful, potentially important information about cochlear and brain health may be missed by thresholds alone. Earlier studies have also incorporated automated or semi-automated peak identification using derivatives (Bradley and Wilson 2004), wavelets (Bradley and Wilson 2005), statistical clustering methods (McCullagh et al. 2007), or a mix of automated and manual techniques (Burke et al. 2023). A few recent studies have used neural network methods similar to those described here, primarily for high sound levels (McKearney et al. 2025, Darahem et al. 2025), as well as across sound levels in normal-hearing and hearing-impaired mice (Erra et al. 2026), which is the closest comparison to the current study. Using our methods, accuracy for animals tested across sound levels and levels of hearing impairment were comparable or exceeded these studies (Fig. 3). These studies suggest convergence upon a successful strategy for rapid ABR analyses that can be extended to other species and conditions. Unlike earlier approaches constrained by explicit morphological assumptions, such as derivative-based latency windows (Fridman 1982; Bradley and Wilson 2005), idealized template matching (Elberling 1979; Vannier 2002; Valderrama 2014), and structural averages in dynamic time warping (Picton 1988; Krumbholz 2020), the CNN learned directly from the data and accommodated local variation in ABR structure. After extrema-based refinement, the large majority of peaks were identified in exact agreement with the ground-truth index rather than a broader tolerance criterion. More permissive scoring schemes can add substantial variability, and even small positional offsets can alter physiological estimates and complicate subsequent interpretation. Minimal adjustments to the initial predictions were sufficient to align them automatically with local extrema, indicating that the model was not reliant on substantial post-hoc correction. Moreover, nearly all adjustments except for wave 5 troughs were < 0.2 ms, and most incorrect adjustments were larger (Fig. 8, Table 5) suggesting that larger adjustments above some set criteria almost certainly indicate guessing or an incorrect prediction by the CNN.

As successful as our classifier was, it can continue to be improved, though additional computations could come with associated increases in processing time. For example, some errors in wave V peaks and troughs appeared to occur due to fusion of two temporally overlapping generators (Kamerer 2024). Because such patterns are challenging for both automated detection and human review, some disagreement likely reflected uncertainty in the reference standard rather than a general limitation of the network. We chose data preprocessing which worked well for our data across conditions, but other settings could be used in cases according to recording configuration and species (Parthasarathy and Kujawa 2018, Laumen et al. 2016, Henry et al. 2011).

## Conclusion

In the present study, ABR response data was obtained by multiple different people recording in a single laboratory and in one species. Although both normal- and altered-hearing recordings were included, the latter was collected from only one exposure paradigm that did not drastically alter waveform timing. Although generalizability across acquisition systems, subject populations, and other etiologies remains to be established, our methods were robust to timing shifts and to below threshold or absent peaks, making them amenable to study noise-induced hearing loss (Henry et al. 2011), blast (Han et al. 2021, Manohar et al. 2020), aging (Sergeyenko et al. 2013, Parthasarathy and Bartlett 2012), or multiple sclerosis (Japaridze et al. 2002), Future work should validate the approach in external datasets spanning additional species and hearing-loss conditions.

## Acknowledgements

The authors thank Devyn Barton, Emily LeBell, Sahil Desai, Audrey Harrison, Amanda Kenney, Hrishikesh Krishna, Valentina Micolisin, Andres Navarro, Elizabeth Schallmo, and Lily Versmesse for assistance with data collection. This work was funded by the Department of Defense (W81XWH2110602 to Edward Bartlett and Aravindakshan Parthasarathy).

## Code availability

Data and code for this project can be found at: https://github.com/jpmarrone/CNN-Rapid-ABR-Peaks.

## Notes

**Conflict of Interest:** The authors declare no competing financial interests.

https://github.com/jpmarrone/CNN-Rapid-ABR-Peaks

### Competing Interest Statement

The authors have declared no competing interest.

